# iPSC Motor Neurons with Familial ALS Mutations Capture Gene Expression Changes in Postmortem Sporadic ALS Motor Neurons

**DOI:** 10.1101/2022.10.25.513780

**Authors:** Aaron Held, Michelle Adler, Christine Marques, Amey S. Kavuturu, Ana R.A.A. Quadros, I. Sandra Ndayambaje, Erika Lara, Michael Ward, Clotilde Lagier-Tourenne, Brian J. Wainger

## Abstract

Motor neuron degeneration, the defining feature of ALS, is a primary example of cell-type specificity in neurodegenerative diseases. Using isogenic pairs of iPSCs harboring different familial ALS mutations, we assess the capacity of iPSC-derived spinal motor neurons, sensory neurons, astrocytes, and superficial cortical neurons to capture disease features including transcriptional and splicing dysregulation observed in human post-mortem neurons. At surprisingly early time points, differentially regulated genes in iPSC-derived spinal motor neurons, but not other cell types, overlap with one-third of the differentially regulated genes in laser-dissected motor neurons from postmortem spinal cords. The extent of dysregulation correlates well between iPSC-derived and *bona fide* spinal motor neurons. In iPSC-derived spinal motor neurons, but not other derived cell types, we detect downregulation of genes affected by TDP-43-dependent aberrant splicing. This reduction takes place exclusively within genotypes known to involve TDP-43 pathology and occurs without evidence of TDP-43 mislocalization or protein level alteration.

## Introduction

Amyotrophic Lateral Sclerosis (ALS) is a fatal neurodegenerative disease characterized by the degeneration of the motor nervous system^1^. Approximately 90% of ALS cases are sporadic (sALS), while mutations in over twenty-five different genes can cause familial ALS (fALS). These fALS mutations sort broadly into functional groups implicated in the disruption of protein homeostasis, RNA processing, or cytoskeletal function^1^, which are exemplified by *SOD1*, *TARDBP*, and *PFN1*, respectively^2^. Nearly all mutations exhibit a dominant inheritance pattern^1^, including the most common genetic cause of ALS, an intronic repeat expansion in the *C9ORF72* gene^3,4^. Two outstanding questions concern the mechanisms underlying selective vulnerability of motor neurons in ALS and convergence of a heterogenous group of genes onto a homogenous clinical presentation.

The degeneration of motor neurons in ALS contrasts with the sparing of other neuronal subtypes^1,5^. Of these, first order sensory neurons, like spinal motor neurons (SMNs), extend long peripheral projections and meet energetic requirements to sustain axonal transport to and from distant compartments. Such similarities between SMNs and sensory neurons juxtaposed with their differential vulnerabilities in ALS provide an opportunity to identify cellular processes that are unique to SMNs and may contribute to their degeneration.

A nearly universal pathological hallmark of ALS is the nuclear depletion and cytoplasmic aggregation of cleaved and phosphorylated TDP-43 (encoded by the *TARDBP* gene), which is found in all sporadic ALS and fALS except fALS due to *SOD1* or *FUS* mutations^6,7^. TDP-43 suppresses human genome-specific cryptic exons in key ALS-related genes^8–12^, leading to an increased emphasis on human cell-based models of ALS, particularly using neurons derived from human induced pluripotent stem cells (iPSCs). However, iPSC-based models of ALS suffer from modest and variable phenotypes, particularly with regard to aspects to of TDP-43 pathology; indeed, many stressors necessary for the emergence of such phenotypes may not be reflective of physiological conditions^13^. Furthermore, because monocultures of iPSC-derived neurons lack critical components of ALS pathogenesis, such as non-cell autonomous toxicity and aging processes^1,14^, mechanisms implicated by studies in iPSC-derived neurons may or may not generalize to less reductionist multi-cellular environments. Critically, there has been little effort to perform systematic validation of candidate disease features identified from iPSC models using primary postmortem tissue, which presumably reflect the integrated contributions of multiple cell types in their native circuit environments.

Here, to address the key questions of cell type specificity and mechanistic convergence, we generated a matrix of different cell types and familial ALS mutations. We used pairs of gene-edited isogenic lines bearing ALS-causing mutations in *SOD1*, *TARDBP*, and *PFN1* and leverage Piggy-Bac transposase to deliver transcription factors for differentiating SMNs, sensory neurons, and superficial cortical-like neurons from each iPSC line. Notably, the effects of ALS mutations varied largely across cell types. SMNs – but not other cell types – captured transcriptional features found in laser-captured SMNs from postmortem ALS spinal cords. Of the three investigated ALS genes, only those known to generate TDP-43 pathology yielded reduced transcripts of genes that undergo TDP-43-dependent aberrant splicing. Finally, in contrast with the detected downregulation of such genes, cryptic exon inclusion events were largely absent in the iPSC SMNs, at least in part reflecting TDP-43 localization patterns and RNA clearance mechanisms that were intact in the iPSC SMNs but become compromised in disease.

## Results

### Generation of different neuronal subtypes using PiggyBac constructs containing inducible transcription factors

Compared to small molecule-based neuronal differentiation, neuronal induction via direct expression of transcription factors in iPSCs can both improve the purity of the target cell population and decrease the differentiation time^15^. To control the expression of transcription factors for differentiation into specific neuronal types, we generated PiggyBac^16^ plasmids with tet-inducible transcription factors to differentiate SMNs (NGN2, ISL1, LXH3)^17^, superficial cortical-like neurons (NGN2)^18^, or first order sensory neurons (NGN2, BRN3A)^19^ (Figure 1A). This approach enables more precise, ratiometric expression of transcription factors than viral transduction strategies while allowing scalability that would be challenging for safe harbor integration strategies^20^. iPSCs with successful integrations were selected using puromycin and then differentiated for 3 days with doxycycline. The resulting neural precursors were either frozen or transferred to long-term media without doxycycline for immediate differentiation and maturation. After 10 days, we used cyclic immunofluorescence^21^ and automated imaging analysis (Figure 1B) to evaluate the SMN markers HB9, CHAT, NEUN, ISL1, and TUJ1. 96.8±0.35% of cells were positive for all markers (Figure 1C, D). We then used RT-qPCR to confirm these expression changes and observed increased *SYN1*, *RBFOX3*, *ISL1*, *MNX1*, and *CHAT* expression in SMNs compared to undifferentiated iPSCs (Figure S1A). A similar analysis of NGN2 superficial cortical-like neurons showed staining for TUJ1 and the layer 2/3 cortical markers BRN2 and FOXG1 (Figure 1E, F); RT-qPCR analysis confirmed increases in the neuronal markers *SYN1* and *RBFOX3* and the upper cortical markers *POU3F2* and *FOXG1*^18^ (Figure S1B). Finally, sensory neurons stained for BRN3A, ISL1, and TUJ1 (Figure 1G, H), and showed increased expression of *SYN1*, *ISL1*, and *POU4F1* by RT-qPCR (Figure S1C).

**Figure 1.**
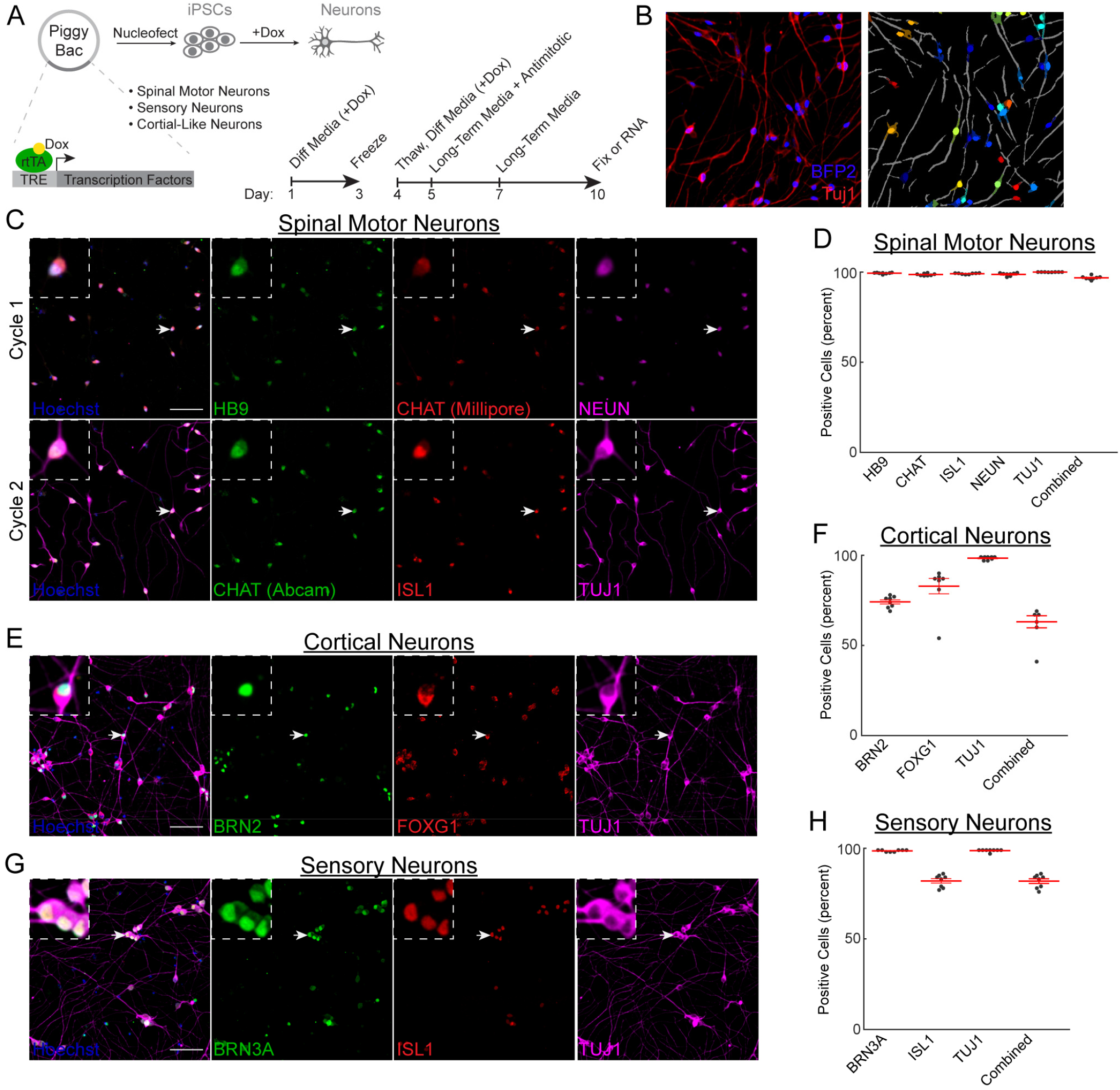
Differentiation of SMNs, cortical neurons, and sensory neurons using inducible transcription factors. (A) Distinct tet-inducible transcription factors contained within PiggyBac constructs were nucleofected into iPSCs to generate different cell types. Timeline summarizes the differentiation protocol. (B) Masking of neurons using BFP2 and TUJ1 staining. Darker, lighter, and gray colors indicate nuclei, cytoplasm, and neurites, respectively. (C) Cyclic immunofluorescence of SMNs for HB9, NEUN, ISL1, TUJ1, and two different CHAT antibodies; quantified in (D). “Combined” indicates the percentage of cells positive for all markers. (E) Cortical neurons stained for BRN2, FOXG1, and TUJ1; quantified (F). (G) Sensory neurons stained for BRN3A, ISL1, and TUJ1; quantified in (H). Scale bars = 100uM; n=8 wells for all groups. Across panels, arrows indicate the same neuron, which is magnified in the inset.

### RNA-sequencing reveals cell-type specific changes resulting from fALS mutations

We next sought to determine which disrupted cellular processes are shared among fALS mutations and whether such perturbations occur specifically in SMNs. Using CRISPR editing, we previously generated isogenic iPSC lines carrying heterozygous fALS mutations (*SOD1^G85R/+^*, *TARDBP^G298S/+^*, and *PFN1^G118V/+^*) and retained unedited iPSC lines that had undergone the editing process as isogenic controls^2^ (Figure 2A). We then generated stable lines carrying inducible transcription factor cassettes and differentiated SMNs and sensory neurons (Figure 1) to identify gene expression changes unique to SMNs as well as those shared among fALS mutations (Figure 2B). We also differentiated astrocytes from all lines^22^ (Figure S2A), as well as NGN2 cortical-like neurons from iPSCs harboring the *TARDBP^G298S/+^* mutation, as about 50% of frontotemporal dementia cases exhibit TDP-43 pathology^6,23^. A principal component analysis (PCA) of all RNA-seq libraries showed a clear separation of astrocytes vs neuronal cell types along principal component 1 (72% of the variance among libraries) as well as a separation of neuronal subtypes along principal component 2 (15% of the variance) (Figure S2B). To confirm the cellular identities of these libraries, we selected cellular markers for our target cell types and performed heatmap clustering (Figure 2C). As expected from the PCA, libraries of the same cell type clustered together and were positive for their respective cell type markers (eg., SMNs with *MNX1* expression, sensory neurons with *SCN9A*, and astrocytes with *GFAP*).

**Figure 2.**
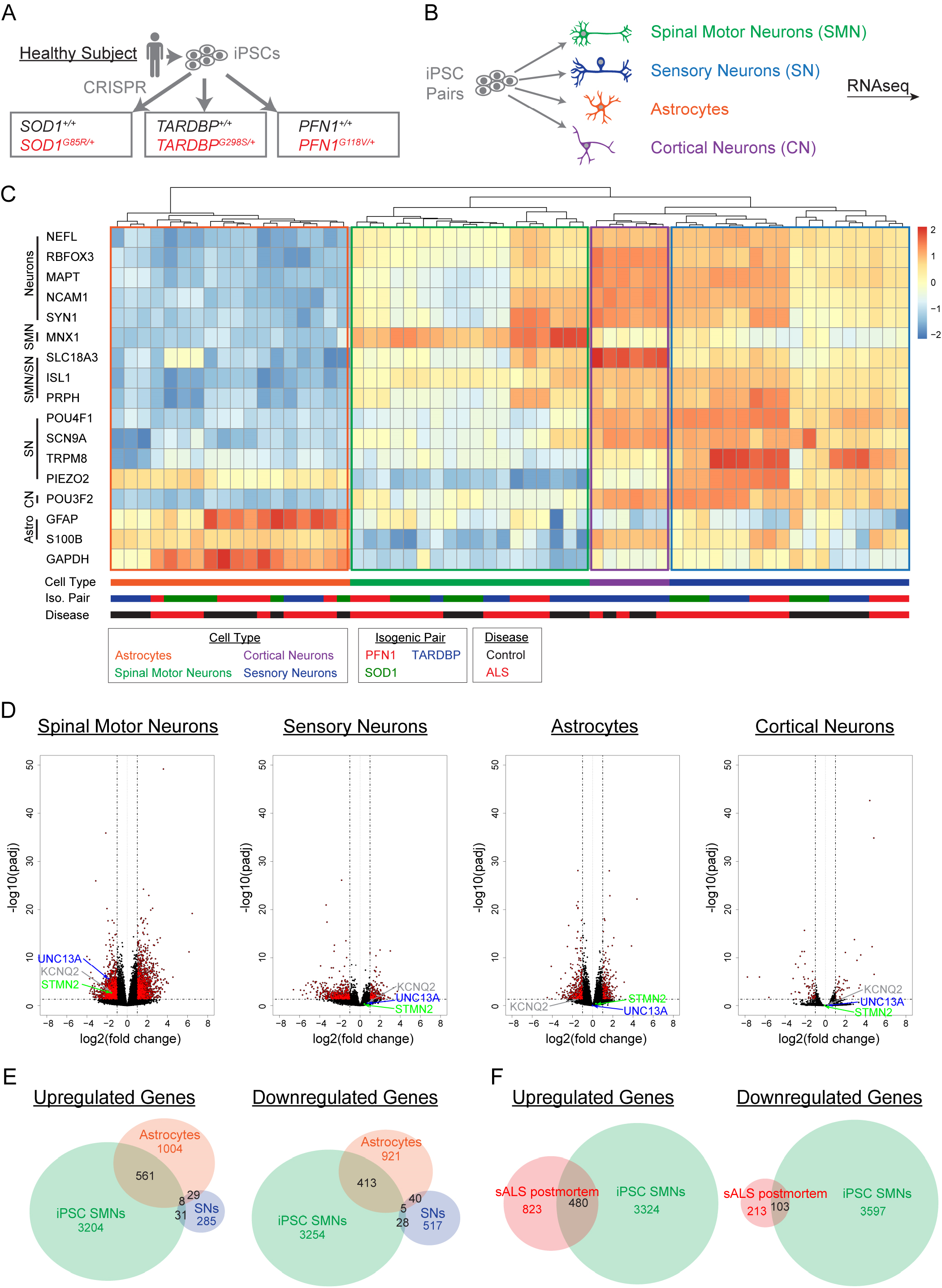
RNA-sequencing of SMNs, sensory neurons, astrocytes, and cortical neurons with fALS mutations. (A) CRISPR editing strategy to generate heterozygous fALS mutations sharing the same genetic background. (B) Differentiation of edited iPSCs into SMNs, sensory neurons, astrocytes, and cortical neurons. (C) Heatmap of all libraries for selected cell markers. (D) Volcano plots comparing all control lines vs all ALS lines across all cell types. (E) Venn diagrams comparing differentially expressed genes due to fALS mutations across different cell types. (F) Venn diagrams comparing differentially expressed genes in fALS iPSC SMNs to differentially expressed genes from laser captured SMNs from postmortem sALS samples (n=8 postmortem control and n=13 postmortem sALS spinal cords).

After verifying the differentiation of our target cell types, we performed an additional principal component analysis within the SMN subgroup to identify differences between SMNs with fALS mutations and controls (Figure S2C). Two of the three SMN genotype isogenic pairs, *TARDBP* and *PFN1*, separated along PC1 (81% of variance), while *SOD1* did not. We then compared iPSC SMNs with fALS mutations against all control libraries and found a substantial overlap in differentially expressed genes across fALS mutations (Figure S2D, Supplementary Table 1). In fact, more genes were shared between all 3 fALS mutations (259 up, 488 down), than were unique to either *SOD1^G85R/+^* (142 up, 70 down) or *PFN1^G118V/+^* (206 up, 358 down). Because of the substantial intersection of differentially regulated genes among all three fALS mutations, we next used a generalized linear model to compare read counts between all control SMNs and all fALS SMNs (Figure 2D). We observed a total of 3,804 upregulated genes and 3,700 downregulated genes between control and fALS SMNs, which was substantially larger than the 354 upregulated and 591 downregulated genes observed when comparing between control and fALS sensory neurons (Figure 2E, Supplementary Table 1). Notably, we observed downregulation of *STMN2*, *UNC13A*, and *KCNQ2*, three genes with important contributions to ALS pathology^9–12,24–26^, in SMNs, but not in sensory neurons, astrocytes, or cortical neurons (Figure 2D). We then compared differentially expressed genes between cell types and observed only a very small overlap between SMNs and sensory neurons, with a more substantial overlap between SMNs and astrocytes (Figure 2E). Gene ontology terms for upregulated genes specific to SMNs included “regulation of programmed cell death” (p=1.7e-9), “cellular response to stress” (p=4.0e-8), and “immune effector process” (p=1.1e-6) while gene ontology terms from downregulated genes specific to SMNs included “modulation of chemical synaptic transmission” (p=6.3e-19), “regulation of neuron projection development” (p=2.6e- 15), and “cell projection organization” (p=2.2e-13) (Figure S2E and Supplementary Table 2).

### iPSC SMNs with fALS mutations capture gene expression changes in postmortem sALS SMNs

To validate the relevance of these expression changes, we compared dysregulated genes in the fALS iPSC SMNs to genes differentially regulated in microdissected SMNs from postmortem sALS spinal cords compared to SMNs from control spinal cords^27^. In the original analysis of this postmortem data set, the top 1000 genes contributing to PC1 were excluded due to concerns regarding microglia contamination. However, 52% of these genes were also differentially expressed in the fALS iPSC SMN cultures, which do not contain microglia (Supplementary Table 3). Thus, an alternative interpretation is that some of these expression changes may genuinely occur within ALS *bona fide* SMNs, consistent with recent findings of traditional inflammatory signaling intrinsic within neurons^28,29^. We then compared our re-analysis of sALS postmortem SMNs (Figure S3A, B) to the fALS iPSC SMNs and found that the fALS iPSC SMNs captured 480/1303 of the upregulated and 103/316 of the downregulated genes in postmortem sALS SMNs (Figure 2F, Supplementary Table 3). Furthermore, the amplitudes of the gene expression changes were also significantly correlated (R=0.74, P=1.2e-102 for intersection genes in Figure 2F, graphed in Figure S3C; R=0.26, p=4.8e-27 for all genes differentially expressed in sALS postmortem SMNs, Supplementary Table 4). Significant GO terms for the overlap between sALS postmortem SMNs and fALS iPSC SMNs included “immune system process” (p=1.7e-15) and “regulation of cell death” (p=5.7e-14) for upregulated genes, and “regulation of cell projection organization” (p=2.2e-5) and “regulation of plasma membrane bounded cell projection” (p=3.6e-5) for downregulated genes (Figure S3D). We then examined differentially expressed genes in postmortem ALS SMNs that were not captured by iPSC SMNs and found “translational initiation” (p=3.5e-13), “cell surface receptor signaling pathway” (p=7.2e-21), and “regulation of response to external stimulus” (p=4.0e-11) in the upregulated genes and no significant GO terms in the downregulated genes (Figure S3E, Supplementary Table 3). Taken together, these data suggest that iPSC SMNs capture a substantial portion of gene expression changes identified in human post-mortem sALS SMNs, while the remaining changes may reflect other contributions such as non-autonomous cellular processes, contributions from aging, and cellular processes specific to sALS.

### Genes with aberrant splicing in ALS patients are downregulated in *TARDBP^G298S/+^* and *PFN1^G118V/+^* SMNs without evidence of mis-splicing

While examining the list of differentially expressed genes specific to SMNs, we noted that several genes shown to exhibit aberrant splicing due to loss of nuclear TDP-43^8–12^, including *STMN2*, *UNC13A*, and *KCNQ2*, were downregulated. In fact, 34/61 genes in which TDP-43-dependent alternative splicing events were observed in FTD-ALS post-mortem neurons^12,30^ were significantly downregulated in the fALS iPSC SMNs (Supplementary Table 5). When we broke down this analysis by isogenic pair, we observed a strong downregulation of these genes in *TARDBP^G298S/+^* and *PFN1^G118V/+^ S*MNs (44/61 and 35/61 genes, respectively), but not in *SOD1^G85R/+^* SMNs (2/61 genes) (Figure 3A). We considered whether this lack of differentially expressed genes in *SOD1^G85R/+^* SMNs may be due to the *SOD1^+/+^* control neurons, since they did not cluster as well with the other control SMNs in our PCA analysis; however, even when comparing *SOD1^G85R/+^* SMNs to all control libraries, only 8/61 genes were differentially expressed. We also did not observe strong downregulation of these genes in sensory neurons or astrocytes for any genotypes (Figure 3A). Thus, the downregulation of these genes results from mutations associated with TDP-43 pathology, *TARDBP^G298S/+^* or *PFN1^G118V/+^* but not *SOD1^G85R/+^*, and occurs specifically in iPSC-derived SMNs.

**Figure 3.**
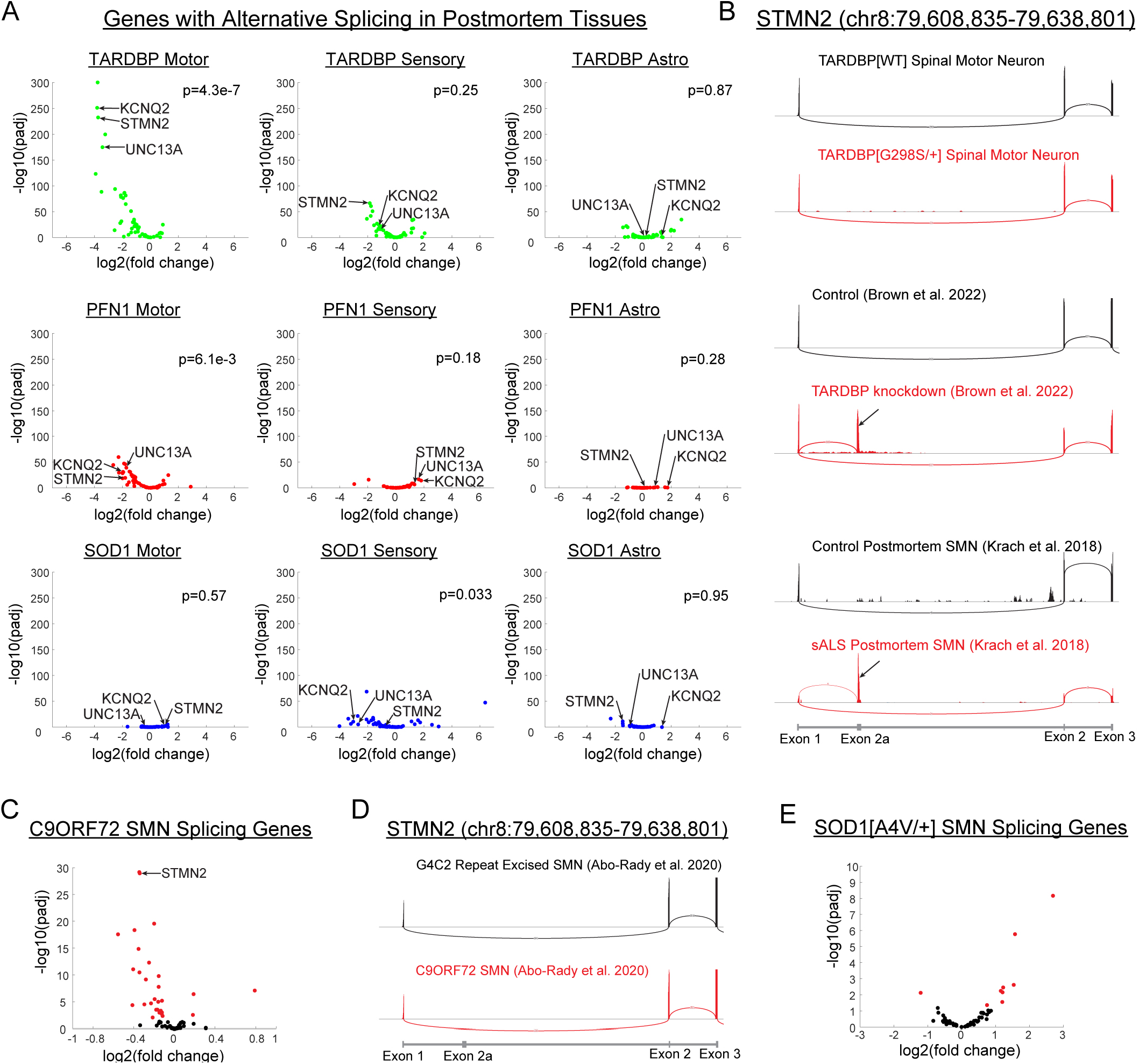
Alternatively spliced genes in TDP-43 depleted FTD-ALS postmortem neurons are downregulated in iPSC SMNs without clear splicing dysregulation. (A) Approximately 80? genes are differentially spliced in postmortem neurons that are depleted for TDP-43 (REF). Those genes are plotted for each cell type and fALS mutation combination. (B) Sashimi plot for *STMN2* splicing in the *TARDBP* SMN isogenic pair, TDP-43 knockdown (Brown et al. 2022), and sALS postmortem SMNs (Krach et al. 2018). Black arrows indicate reads aligning to exon 2a. (C) Expression of splicing genes in *C9ORF72* iPSC SMNs (Abo-Rady et al. 2020), significant genes colored in red; n=6 libraries per condition. (D) Sashimi plot of *STMN2* splicing in *C9ORF72* iPSC SMNs. (E) Expression of splicing genes in *SOD1^A4V/+^* iPSC SMNs (Kiskinis et al. 2014), significant genes colored in red; n=3 control libraries and n=2 *SOD1^A4V/+^* libraries.

We next considered whether these genes are downregulated because iPSC SMNs recapitulate the aberrant splicing observed in FTD-ALS post-mortem neurons with depleted nuclear TDP-43^12,30^. Some of these alternative splicing events, such as splicing in cryptic exons with early stop codons, yield transcripts that are cleared by nonsense mediated decay (NMD) or lead to premature polyadenylation^9–11^. As a control, we re-analyzed four published data sets in which cryptic splicing events have been reported^10–12^ and were able to replicate detection of cryptic exons (Figure 3B, Figure S4A, B, Supplementary Table 5). Although we observed clear inclusion of the *STMN2* cryptic exon and several other cryptic splicing events in these data sets, we did not observe any cryptic exon inclusion in the fALS iPSC SMNs. We were able to replicate seven alternative splicing events identified in post-mortem FTD-ALS neurons in fALS iPSC SMNs (Supplementary Table 5), but none of these events cause cryptic exon inclusion. The apparent dichotomy between the downregulation of these genes and the lack of cryptic exon detection led us to evaluate additional previously published datasets of *C9ORF72* and *SOD1^A4V/+^* iPSC SMNs^31,32^ and examine the same subset of genes^12,30^. In *C9ORF72* iPSC SMNs^32^, we found that 26/61 genes were significantly downregulated (Figure 3C) again without any cryptic splicing events in these genes (Figure 3D, Figure S3). Only one gene was decreased in *SOD1^A4V/+^* iPSC SMNs^31^ (Figure 3E), consistent with our own results in *SOD1^G85R/+^* iPSC SMNs (Figure 3A).

To account for the possibility of RNA-sequencing artifacts preventing detection of cryptic exons in the iPSC SMNs, we performed RT-qPCR against the *STMN2* and *UNC13A* full-length genes and their cryptic exons (Figure 4A, Figure S4C). The expression of the full-length genes mirrored the RNA-sequencing results, and we did not observe any increase in cryptic exon expression. We obtained similar results by performing RT-qPCR with hydrolysis probes (Figure S4D). We then quantified TDP-43 protein expression and localization in iPSC SMNs and found no change in TDP-43 expression and only minimal difference in localization (Figure 4B, C, Figure S4E, S4F).

**Figure 4.**
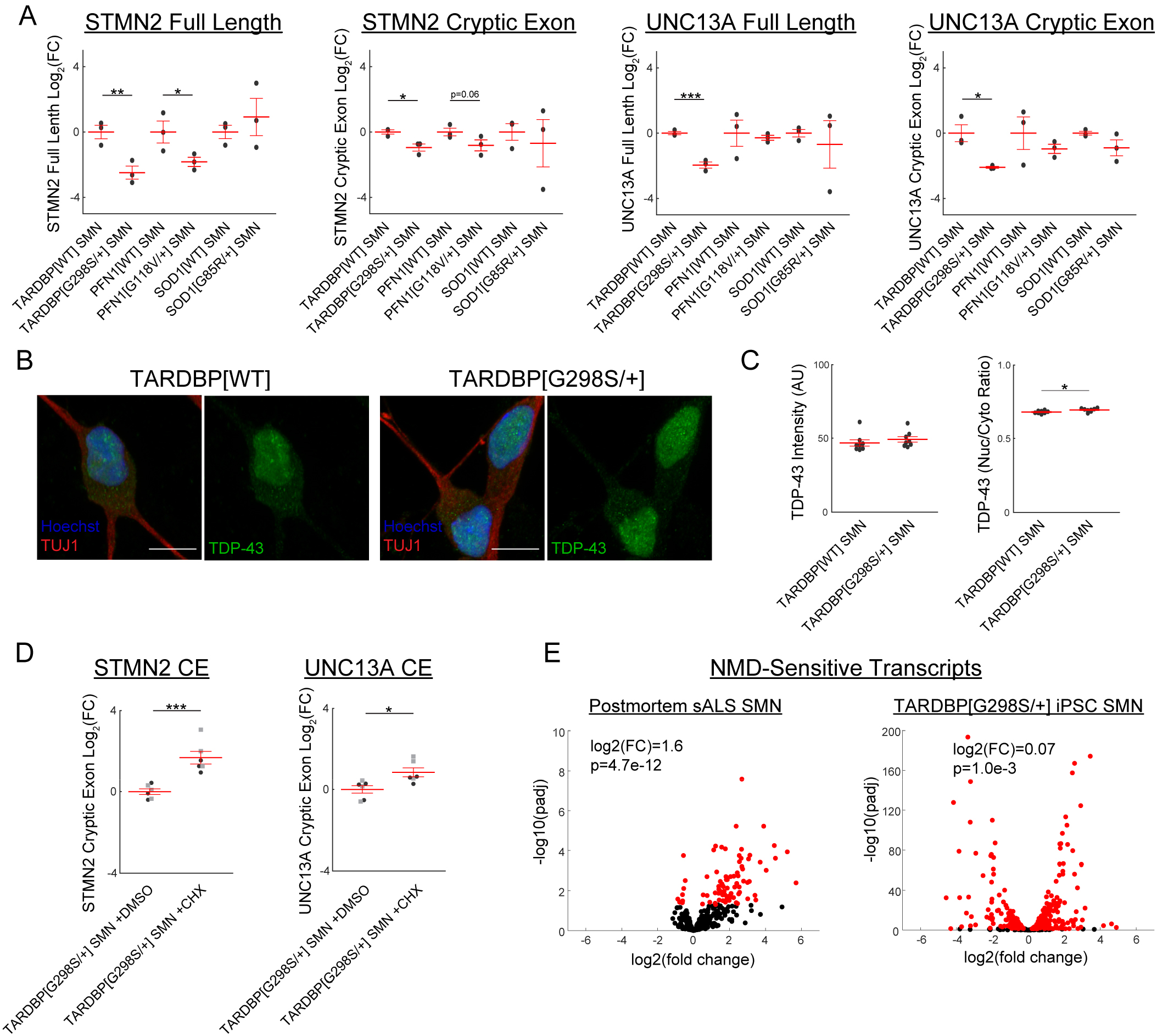
Transcripts containing cryptic exons are rapidly degraded. (A) RT-qPCR for full length transcripts and cryptic exon transcripts in *STMN2* and *UNC13A*; n=3 wells per condition. (B) TDP-43 protein expression and localization in *TARDBP^G298S/+^* and *TARDBP^+/+^* iPSC SMNs (scale bar = 10um); quantified in (C); n=8 wells per genotype. (D) RT-qPCR for cryptic exon transcripts after inhibiting translation with CHX. Two separate experiments were conducted, one denotated by black circles and the other by gray squares. (E) Expression of NMD-sensitive transcripts in postmortem sALS SMNs and *TARDBP^G298S/+^* iPSC SMNs.

### Cryptic splicing in *TARDBP^G298S/+^* SMNs is masked by rapid transcript degradation

The lack of cryptic exons in the *TARDBP^G298S/+^* and *PFN1^G118V/+^* SMNs and previously published heterozygous *C9ORF72* iPSC SMNs contrasts with the detection of cryptic exons in previous publications using homozygous *TARDBP* mutations and *TARDBP* knockdown^9–11,33^. Notably, homozygous *TARDBP* mutations led to less frequent cryptic exon inclusion than *TARDBP* knockdown (Figure S4A, B), leading us to hypothesize that cryptic exon inclusion may occur at an even lower rate with only heterozygous *TARDBP* mutations. In general, aberrant mRNA transcripts are degraded by translation-dependent RNA clearance mechanisms^34^, including the nonsense-mediated decay (NMD) pathway but also no-go and non-stop mechanisms^11^. To determine whether RNA clearance mechanisms degraded cryptic exon-containing transcripts, we treated *TARDBP^G298S/+^* iPSC SMNs with cycloheximide for six hours to block translation-dependent RNA clearance and then performed RT-qPCR for *STMN2* and *UNC13A* cryptic exons. We detected an increase in both *STMN2* and *UNC13A* cryptic exons after CHX treatment (Figure 4D), indicating that these transcripts accumulate quickly when translation-dependent RNA quality control measures are blocked.

The rapid clearance of transcripts containing cryptic exons led us to hypothesize that RNA clearance mechanisms may be functional in iPSC SMNs, but disrupted in sALS postmortem SMNs, and thus impairment of RNA clearance may lead to cryptic exon accumulation in postmortem SMNs. To test this possibility, we used upregulation of established NMD target genes as an index of blocked NMD^35–37^ and probed NMD integrity in postmortem sALS SMNs and the fALS iPSC SMNs. We observed an increase in the expression of NMD-sensitive transcripts in postmortem sALS SMNs (log2(Fold Change)=1.6), indicating a substantial deficit in NMD (Figure 4E). In contrast, we observed much smaller effects on NMD target genes in fALS iPSC SMNs (*TARDBP^G298S/+^* = 0.07, *PFN1^G118V/+^* = 0.49, *SOD1^G85R/+^* =0.08) (Figure 4E, S4G). Thus, intact RNA quality control mechanisms may eliminate cryptic exon transcripts in fALS iPSC SMNs whereas impairment of such mechanisms in postmortem neurons may contribute to cryptic exon persistence and consequent detection at typical RNA-sequencing read depths.

## Discussion

iPSC-based modeling studies of neurodegenerative disease have implicated individual candidate genes that have been validated in postmortem disease tissue^9–13,38^. However, there have not been systemic comparisons to address, on a genome wide level, how comprehensively iPSC models capture gene dysregulation in *bona fide* disease-relevant cell types. We find that after only 10 days in culture, SMN monocultures capture a full third of the genes differentially regulated in sALS laser-captured SMNs from post-mortem tissue compared to control. The correlation of effect size for genes within the intersection of the iPSC and postmortem SMN pools is striking, with R=0.74, although lower but still strongly significant when genes outside the intersection are included. This result broadly validates iPSC modeling using disease-relevant cell types and helps delineate which effects are captured in monocultures and which result from non-autonomous or aging effects. Furthermore, such systematic comparisons of iPSC models and postmortem tissue may provide reciprocal insight into the interpretation of the postmortem datasets. For example, in the iPSC SMNs, which do not contain immune cells, the dysregulation of genes typically associated with innate immune activation may support greater impact of such signaling pathways within neurons themselves^28^, and less influence of microglial contamination in laser-captured postmortem SMNs^27^.

By using a matrix of iPSCs harboring different familial ALS mutations and a range of differentiated cell types, we address both how accurate disease modeling is in the most vulnerable cell type, namely SMNs, and how specific are the disease features across the different cell types. We initially hypothesized that fALS mutations would cause similar gene expression changes in first order sensory neurons and SMNs because of the shared morphological and consequent energetic and trafficking challenges, and therefore the differences would provide insight into the selective vulnerability of SMNs in ALS, as have been seen by comparing vulnerable and less vulnerable motor neurons^39,40^. However, we observed almost completely different sets of differentially expressed genes in SMNs and sensory neurons, illustrating the extent to which ALS mutations initiate broadly different transcriptional programs in vulnerable versus non-vulnerable cell types. Similar to findings for the sensory neurons, the *TARDBP^G298S/+^* mutation resulted in markedly different gene expression changes in superficial cortical-like NGN2 neurons compared to SMNs. These results emphasize how cell fate dramatically influences downstream responses, even between similar cell types, and highlights the necessity of using the most relevant cell type when modeling neurodegenerative diseases. Within the SMNs, the substantial overlap among dysregulated genes across all three ALS genotypes supports a mechanism of phenotypic convergence in ALS and provides candidate genes that may be subsequently tested for such a role. Given the demonstrated value of disease modeling with specific cell types and multiple genotypes, the capacity of PiggyBac integrations to provide robust, rapid, and highly pure transcription factor-based differentiations across a range of iPSC lines and cell types will be useful for accurate modeling of selective vulnerability in neurodegenerative disease at robust scale.

The example of transcript depletion of genes that require TDP-43 for proper splicing illustrates the specificity of iPSC modeling for both cell type- and genotype-specific effects^8–12,30^. While TDP-43 depletion or homozygous mutation of TDP-43 elicited aberrant splicing in multiple cell types^8–12,30,33^, we found that heterozygous mutations – as occur in the disease – produced downregulation of such transcripts in SMNs, but not in sensory neurons, NGN2 neurons, or astrocytes. This interaction between manifestation of TDP-43 dysfunction and cellular identity is also consistent with previous reports of *TARDBP* knockdown yielding different cryptic splicing events in embryonic stem cells and NGN2 neurons^8,11^. In addition to cell type effects, the transcript abundance of these genes show genotype-specific patterns that parallel the presence or absence of TDP-43 pathology in human tissue: the decrease in transcript abundance occurred in fALS iPSC SMNs harboring *TARDBP*, *PFN1*, and *C9ORF72* mutations, genes linked to TDP-43 pathology, but not in two different *SOD1* mutations, which do not yield appreciable TDP-43 pathology in postmortem tissue^6,7^.

Recent models of TDP-43 posit that loss of nuclear TDP-43 or TDP-43 dysfunction produces a decrease in full length transcripts of TDP-43-depedent splicing genes as well as concomitant accumulation of cryptic exons due to aberrant misplicing.^8–12,30^. While we observed a reduction in the abundance of these full-length transcripts in iPSC SMNs harboring *TARDBP* or *PFN1* mutations, we did not detect cryptic exon transcripts directly in our RNA-sequencing data or in datasets from similar studies^31,32^, suggesting that cryptic exon transcripts were either absent or present only in low abundance. Thus, our findings yielded a dichotomy between the detected decrease in full length transcripts and the lack of expected reciprocal increases in cryptic ones. To explain this surprising finding, we hypothesized that transcripts containing cryptic exons could be degraded in the iPSC SMNs by one of many translation-dependent RNA clearance pathways such as NMD, no-go, and non-stop mechanisms^11,34^. Broadly blocking such RNA clearance mechanisms in *TARDBP^G298S/+^* SMNs with cycloheximide yielded an increase in cryptic exon transcripts for *STMN2* and *UNC13A*, consistent with the interpretation that intact protein synthesis-dependent RNA clearance may sharply decrease cryptic exons in iPSC SMNs. These results differ slightly from experiments performed in SH-SY5Y cells, in which blocking translation-dependent RNA clearance increased cryptic exon abundance for *UNC13A* but not *STMN2*^11^. Nonetheless, given our demonstration of strong cell type influence on TDP-43-related aberrant splicing, it would not be surprising if RNA clearance mechanisms differed between SH-SY5Y cells and iPSC SMNs as well. Consistent with such cell type differences in RNA clearance, analysis of consensus NMD decay target levels did not find NMD impairment in sALS brain tissue^41^, but our similar analysis of postmortem sALS SMNs^27^ strongly suggests NMD is impaired. Notably, the impaired NMD in the sALS SMNs contrasted with the intact NMD index in the iPSCs SMNs. Thus, under conditions of physiological TDP-43 perturbation, dysfunction in RNA clearance mechanisms in postmortem SMNs may contribute to the persistence of cryptic exon transcripts while intact clearance mechanisms degrade cryptic exons in iPSC SMNs.

The excitement and rapid advancement in TDP-43-dependent splicing has left many questions unanswered, including impact of cell type and magnitude of TDP-43 depletion, for which our results provide some insight, as well as interactions between these factors and specific cryptic splicing events for different genes. Our results do not further resolve the specific translation-dependent RNA mechanisms that clear cryptic exon transcripts, because these likely vary between specific transcripts and because the basic biology of these mechanisms and the capacity to pharmacologically manipulate them are both limited. Similarly, additional influence of greater cellular age or addition of non-cell autonomous features may affect RNA clearance and consequently cryptic exon detection in SMNs. Nonetheless, because of the apparent sensitivity of cryptic detection to RNA clearance, emphasis on depletion of full-length transcript levels may prove a valuable focus for cellular models and biomarker development.

## Methods

### iPSC culture

iPSCs were plated on Vitronectin XF (Stemcell Technologies 07180) using mTeSR Plus (Stemcell Technologies 100-0275) supplemented with CET^42^. After day 1, CET was removed, and cells were maintained in mTeSR Plus. Cells were passaged with accutase (Thermofisher Scientific A1110501). All iPSC lines have been described previously^2^.

### Plasmids

All donor plasmids and their sequences will be made publicly available through Addgene at the time of publication.

### Nucleofection

iPSCs were nucleofected using the Human Stem Cell Nucleofector Kit 1 (Lonza VPH-5012) and protocol A- 23 with a Nucleofector I (Lonza). Both the transposase plasmid (2.75ug) and donor plasmid (2.75ug) were nucleofected into 800,000 iPSCs and then maintained in mTeSR Plus with CET for 24 hours. Cells with successful nucleofection were then selected using puromycin (10ug/mL, InvivoGen ant-pr-1) for 48 hours. Selection continued until all remaining cells had a nuclear BFP2 signal, indicating successful integration of the donor plasmid.

### Neuron Differentiation

iPSCs were differentiated using induction media (480mL DMEM/F12, Life Technlologies 11320082; 5mL N2 supplement, Gibco 17502-048; 5mL non-essential amino acids, Corning 25-025-CI; 5mL Glutamax, Thermo 35050061; 5mL Penicillin/Streptomycin, Life Technologies 15070-063) supplemented with doxycycline (2ug/mL, Sigma Aldrich D9891-1G) and CET. For SMN differentiations, Compound E (0.2uM, Calbiochem 565790-500UG) was also added. 10 million iPSCs with integrated donor plasmids were resuspended in 25mL supplemented induction media and plated in a T175 flask coated with Matrigel (Corning 354277). After 24 hours, the media was exchanged for fresh supplemented induction media. The following day differentiated cells were either frozen in batches of 1 million cells/vial (250uL supplemented induction media, 200uL FBS (Hyclone SH30910.03HI), 50uL DMSO (Sigma D2650)) or replated for experiments.

For immunostaining experiments, 96-wells plates (Cellvis P96-1.5H-N) were coated with PEI (Sigma 03880) diluted 1:50 in borate buffer (100mM boric acid, 25mM sodium tetraborate, 75 mM sodium chloride, pH 8.4) for 24 hours and then washed 5x in 1xPBS (Gibco 10010049). They were then coated for an additional 24 hours in laminin (12ug/mL, Life Technologies 23017-015) and then washed an additional 3x in 1xPBS. Frozen, differentiated neurons were thawed in supplemented induction media and plated at either 40,000 cells/well (Figure 1) or 80,000 cells/well (Figure 3). The following day, neurons were transferred to long-term media supplemented with aphidicolin (5uM, Cell Signaling Technology 32774). After two days, media was exchanged for fresh long-term media without aphidicolin. Neurons were cultured for an additional 3 days and then fixed for immunofluorescence. For RT-qPCR experiments, frozen cells were thawed into 6-well plates (Corning 353046) coated in Matrigel at a density of 1 million cells/well in supplemented induction media. The next day, media was exchanged for long-term media with aphidicolin. Two days later, media was exchanged for long-term media without aphidicolin. Cells were harvested 3 days later by direct application of buffer RLT from the RNeasy mini kit (Qiagen 74104).

SMN long-term media consisted of Neurobasal (478mL, Life Technologies 21103049), N2 supplement (5mL, Gibco 17502-048), non-essential amino acids (5mL, Corning 25-025-CI), Glutamax (5mL, Thermo 35050061), Penicillin/Streptomycin (5mL, Life Technologies 15070-063), and beta mercaptoethanol (2mL, Sigma M6250-100ML). SMN media was supplemented with GDNF (10ng/mL, Life Technologies PHC7044), BDNF (10ng/mL, Life Technologies PHC7074), CTNF (10ng/mL R&D Systems 257-NT-50UG), IGF-1 (10ng/mL, R&D Systems 291-G1-200), and retinoic acid (1uM, Sigma R2625-50MG).

Long-term media for cortical-like neurons and sensory neurons consisted of Neurobasal (475mL, Life Technologies 21103049), B27 supplement (10mL, Gibco 17504-44), Glutamax (5mL, Thermo 35050061), Penicillin/Streptomycin (5mL, Life Technologies 15070-063), and sodium chloride (5mL of 5M solution, Sigma 71376-1KG). Cortical neuron media was supplemented with NT-3 (10ng/mL, PeproTech, 450-03) and BDNF (10ng/mL, Life Technologies PHC7074). Sensory neuron media was supplemented with NT-3 (10ng/mL, PeproTech, 450-03), BDNF (10ng/mL, Life Technologies PHC7074), GDNF (10ng/mL, Life Technologies PHC7044), and NGF (10ng/mL, R&D Systems 256-GF).

### Astrocyte Differentiation

1 million iPSCs were resuspended in 12mL N2B27 media (250mL Advanced DMEM/F12, Thermo 12634-028; 250mL Neurobasal, Life Technologies 21103049; 5mL Penicillin/Streptomycin, Life Technologies 15070-063; 5mL Glutamax, Life Technologies 35050061; 1mL beta mercaptoethanol, Sigma M6250-100ML; 10mL B27, Life Technologies 17504044; 5mL N2 Life Technologies 17502048) supplemented with LDN193189 (0.1uM, Stemgent 04- 0074-02), SB431542 (20uM, DNSK DNSK-KI-12), Retinoic Acid (100nm, Sigma R2625-50MG), and Y27632 (10uM, abcam ab120129), and plated in a T75 flask coated with Matrigel. The following day, fresh supplemented N2B27 media was exchanged, except without Y27632. Media was exchanged again on day 4. On day 6, neural stem cells were lifted with accutase and 1 million cells were resuspended in 12mL astrocyte media with astrocyte supplement, FBS, and Penicillin/Streptomycin (all astrocyte media reagents included in Sciencell 1801) and then plated in a T75 flask coated with Matrigel. Media was exchanged every 2-3 days for 1 month. When cells reached 80% confluence, they were split 1:4 into a new T75 flask coated with Matrigel.

For immunofluorescence experiments, astrocytes were seeded into 96-wells plates (Cellvis P96-1.5H-N) coated with PEI and laminin (see neuron differentiation). 10,000 astrocytes per well were seeded and then grown for an additional week in supplemented astrocyte media before fixation with formaldehyde (Life Technologies #28908).

### SH-SY5Y Culture

The neuroblastoma SH-SY5Y cells (ATCC) were cultured in DMEM F12 (Gibco 1132-033) containing 1% of Penicillin-Streptomycin (Gibco 15140-122) and 10% of Fetal Bovine Serum (Sigma F0926) and kept at 37oC and 5% CO2. The knockdown was achieved by transfecting cells for 48h with siRNA against TDP-43 (ON-TARGETplus siRNA, Dharmacon L-012394) or a control sequence (ON-TARGETplus Non-targeting Control Pool, Dharmacon D001810–10) using Lipofectamine RNAiMAX Transfection Reagent (13778075) in Opti-MEMTM (Gibco 31985070). We used a final siRNA amount of 2.5pmol per well.

### Cycloheximide Treatment

iPSC SMNs were grown to day 10 as described in the neuron differentiation methods. iPSC SMNs were treated with DMSO or 100uM Cycloheximide (Millipore Sigma 01810) for 6 hours and then harvested using buffer RLT from the RNeasy mini kit (Qiagen 74104).

### Immunofluorescence

Cells were fixed using using 4% formaldehyde (Life Technologies #28908) for 15 minutes and then blocked and permeabilized for 1 hour in PBT (1xPBS, Gibco #10010049; 0.5% Triton-X, Millipore Sigma #9400; 1% BSA, Millipore Sigma #A2153). Cells were then incubated overnight in PBT with primary antibodies. The following day, cells were washed 2x in 1xPBS and incubated for 1 hour in PBT with secondary antibodies and Hoechst (1:1000, Thermofisher Scientific 62249). Cells were then washed 2x in 1xPBS and then imaged with an ImageXpress micro confocal (MetaXpress version 6 software; 10x objective) or a Zeiss LSM900 (Zen 3.3 software; 20x or 63x objective). Images were quantified using custom MATLAB (2018b) scripts (https://github.com/waingerlab/iPSC-SMN). Briefly, the nuclear signal was thresholded at 3 standard deviations above the mean of the background signal, and groups of pixels meeting this threshold were identified as nuclei. The same approach was used with TUJ1 to identify the cytoplasmic compartment. Thin segments of the TUJ1 signal extending away from the cytoplasm were classified as neurites. A cell was considered “positive” for a marker if the average of the cell’s signal for that marker was more than 2 standard deviations above the background noise.

The primary antibodies used in this study were: anti-Brn2 (1:1000, Cell Signaling 12137S), anti-FoxG1 (1:500, Millipore Sigma MABD79), anti-Tuj1 (1:500, Aves TUJ-0020), anti-Brn3a (1:200, Millipore MAB1585), anti-Isl1 (1:200, Abcam ab203406), anti-GFAP (1:250, abcam ab194324), anti-S100β (1:200, abcam ab52642), and anti-TDP-43 (1:500, proteintech 10782-2-AP). Secondary antibodies used in this study were: donkey anti-mouse 488 (1:1000, Thermofisher Scientific A21202), donkey anti-mouse 568 (1:1000, Thermofisher Scientific A10037), donkey anti-rabbit 488 (1:1000, Thermofisher Scientific A21206), donkey anti-rabbit 568 (1:1000, Thermofisher Scientific A10042), and donkey anti-chicken 647 (1:1000, Jackson Immunoresearch 703-605-155).

### Cyclic Immunofluorescence

SMNs were first imaged before immunostaining to track the BFP2 signal. They were then blocked and permeabilized for 1 hour in PBT and incubated overnight in a primary antibody solution of: PBT, anti-Hb9 (1:150, DSHB 81.5C10), anti-CHAT (1:100, EMD millipore AB144P-200UL), and anti-NeuN (1:500, EMD millipore ABN78). Cells were then washed 2x in 1xPBS and incubated for 1 hour in a secondary solution of: PBT, Hoechst (1:1000, Thermofisher Scientific 62249), donkey anti-mouse 488 (1:1000, Thermofisher Scientific A21202), donkey anti-goat 555 (1:1000, Thermofisher Scientific A21432), and donkey anti-rabbit 647 (1:1000 Thermofisher Scientific A31573). Cells were then imaged on an ImageXpress micro confocal. Fluorophores were then bleached overnight using 25mM sodium hydroxide (Sigma-Aldrich 72068-100ML) in 1xPBS with 4.5% hydrogen peroxide (Thermofisher Scientific H325-500). Fluorophore bleaching was confirmed using an ImageXpress micro confocal before proceeding to additional staining. Cells were then stained overnight with a primary solution of: PBT, anti-CHAT 488 (1:100, ab192465), anti-Isl1 568 (1:200, ab203406), and anti-Tuj1 (1:500, Aves TUJ-0020). They were then washed 2x in 1xPBS and incubated for 1 hour in a secondary solution of PBT with donkey anti-chicken 647 (Jackson Immunoresearch 703-605-155) and Hoechst (1:1000, Thermofisher Scientific 62249). Cells were washed an additional 2x in 1xPBS and imaged on an ImageXpress micro confocal. Images were then aligned between cycles using custom MATLAB scripts (https://github.com/waingerlab/iPSC-SMN).

### RNA extraction and RT-qPCR

Cells were harvested using buffer RLT from the RNeasy mini kit (Qiagen 74104). The RNeasy mini kit was then used to extract RNA, with QIAshredder columns (Qiagen 79654) used to increase the yield. The High-Capacity cDNA Reverse Transcription Kit (Applied Biosystems 4368814) was used to make cDNA. KiCqStart SYBR Green (Millipore Sigma KCQS00) was used with a BioRad CFX96 PCR system (running BioRad CFX Manager 3.1) to perform RT-RT-qPCR. Primers are listed in Supplementary Table 6. All gene expression was normalized to *GAPDH*, and fold change was calculated using ΔΔCt.

For RT-qPCR with hydrolysis probes, the *GAPDH* primers were: GAAGGTGAAGGTCGGAGTC (forward), GAAGATGGTGATGGGATTTC (reverse), and CAAGCTTCCCGTTCTCAGCC (probe). The *STMN2* cryptic exon primers were CTTTCTCTAGCACGGTCCCAC (forward), ATGCTCACACAGAGAGCCAAATTC (reverse), and CTCTCGAAGGTCTTCTGCCG (probe). cDNA, primers, and iTaq (BioRad 1725131) were mixed together and analyzed using a BioRad CFX96 PCR system.

### RNA sequencing

Separate differentiations were completed for each replicate. 1 million cells per differentiation were pelleted and sent to Genewiz for RNA extraction, library preparation (TruSeq RNA Library Prep Kit v2, mRNA selection), and RNA sequencing (Illumina HiSeq). An average of 26.5 million 2×150bp paired end reads were obtained per library. Reads were aligned to GRCh38 using STAR (2.7.3a) with refgene annotation and counted using STAR --quantmode GeneCounts. Differential expression was calculated using DESeq2 (1.36.0) in R (4.0.3). To identify general differences between ALS and control libraries, we used design = ~Gene+Disease, where “Gene” indicates the isogenic pair and “Disease” represents whether a mutation that causes ALS is present. PCA plots, heatmaps, and volcano plots were made in R (4.0.3) or MATLAB (2018b), Venn Diagrams in MATLAB (2018b), and Sashimi plots in IGV (2.11.9). For differential splicing analysis, reads were aligned to GRCh38 using STAR (2.7.3a) with Gencode v40 annotation. Aligned bams were indexed with samtools (1.13) and differential splicing identified with Majiq (2.3). Reads aligning to cryptic exons were counted using bedtools (2.27.1). The previously published datasets reanalyzed in this study were obtained through the NCBI GEO database (GSE54409^31^, GSE143743^32^, GSE76220^27^, GSE122069^10^, GSE126543^30^) or the European Nucleotide Archive (PRJEB42763^11^). For the reanalysis of postmortem sporadic ALS SMNs^27^, a model of design=~Subject_Sex+Disease was used to normalize for effects by patient sex. For the reanalysis of TDP-43 sorted neurons^30^, a model of design =~Subject+TDP-43_status was used to pair libraries within the same patient. Genes with FDR<0.05 were considered significantly differentially expressed and used in downstream analyses. Differentially expressed genes were analyzed for gene ontology using GOrilla^43^ and summarized with Revigo^44^. Only GO terms with corrected p-values less than 0.001 were considered significant. For the analysis of differential splicing events described in Ma et al. 2022, 5 genes were excluded from our analysis because either there were not enough reads in our RNA-seq libraries to evaluate differential expression or the gene symbols did not match because of the use of different annotation files between studies.

### NMD-Sensitive Transcripts

Previous studies identified differentially expressed genes after UPF1^35^, UPF2^36^, or UPF3^37^ depletion. We compiled the upregulated genes (Supplemental Table 7) as a readout NMD function and then assessed the expression of these transcripts in iPSC SMNs and postmortem SMNs. Significance was calculated using fgsea (1.22.0) in R (4.0.3), and the average log2(fold change) was determined by averaging the log2(fold change) of differentially regulated genes (adjusted p<0.05).

### Statistical analysis

For RNA sequencing experiments, each replicate was obtained from a separate batch differentiation. A negative binomial generalized linear model (DESeq2 1.36.0) was used to identify differences in gene expression. For immunofluorescence and RT-qPCR experiments, neurons were derived from the same batch differentiation, frozen into separate vials, and then plated in either 96-well or 6-well format. Individual plotted dots represent the average of technical replicates in a well (RT-qPCR) or the average of individual cells in a well (immunofluorescence). Red bars represent the average and standard error of the mean. Two sided t-tests (MATLAB 2018b) were used to compare between groups. Pearson correlations (MATLAB 2018b) compared differentially expressed genes in postmortem sALS SMN to differentially expressed genes in iPSC-neurons. Significance for downregulated alternative splicing genes was calculated with fgsea (1.22.0) in R (4.0.3). P-values are depicted in graphs as *, **, or ***, representing p<0.05, p<0.01, and p<0.001, respectively.

## Data availability

RNA-sequencing data generated in this manuscript will be made publicly available at the time of publication. Other raw data and CRISPR edited iPSC lines will be made available upon reasonable request to the authors.

## Code availability

Custom scripts used to analyze immunofluorescence and RNA-seq data are available at https://github.com/waingerlab/iPSC-SMN.

---

**Figure S1.**
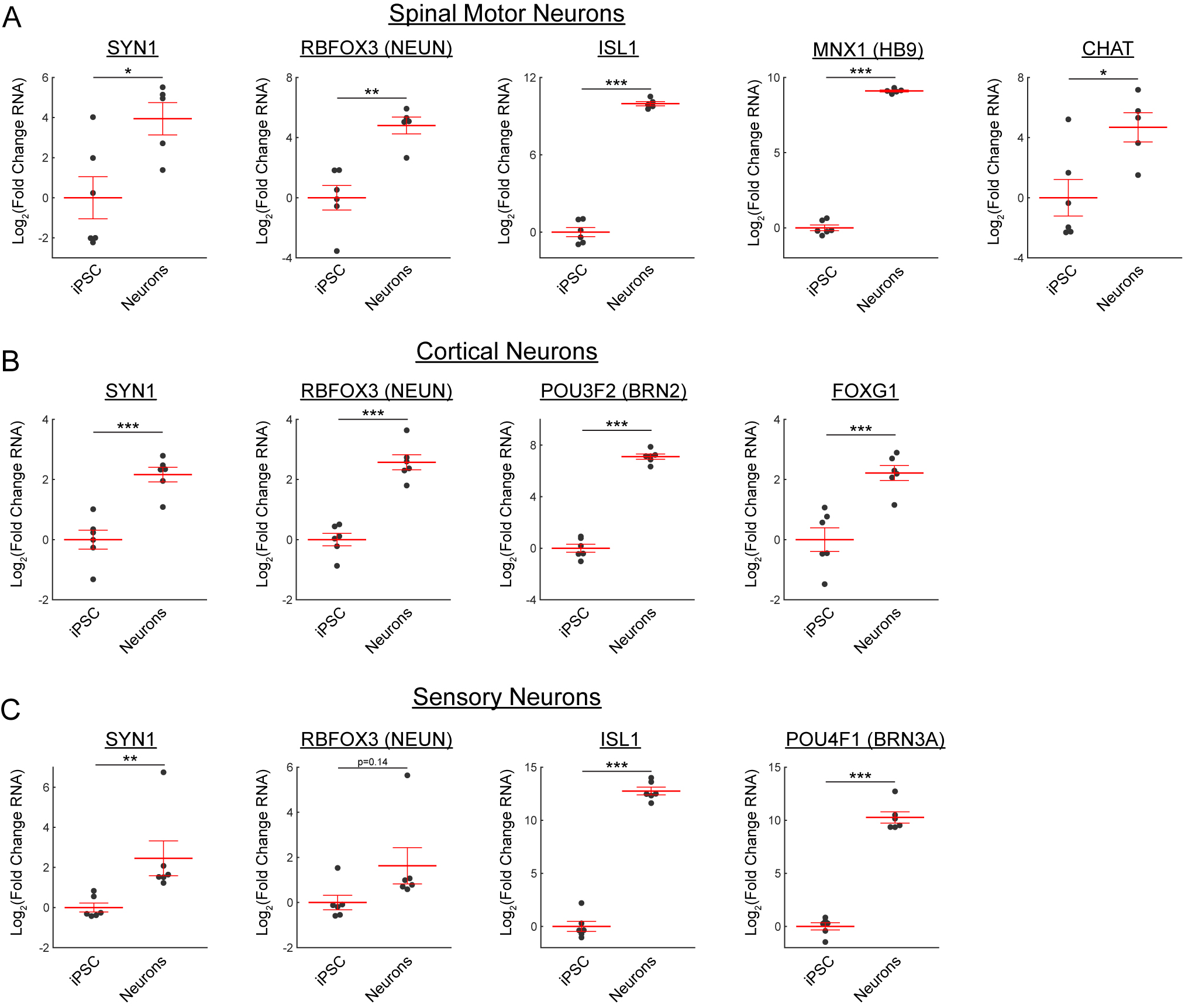
RT-qPCR analysis for SMN, cortical neuron, and sensory neuron markers. (A) SMN markers in SMNs, (B) cortical neuron markers in NGN2 neurons, and (C) sensory neuron markers in sensory neurons; normalized to *GAPDH* and undifferentiated iPSCs containing the differentiation construct; n=6 wells for all groups.

**Figure S2.**
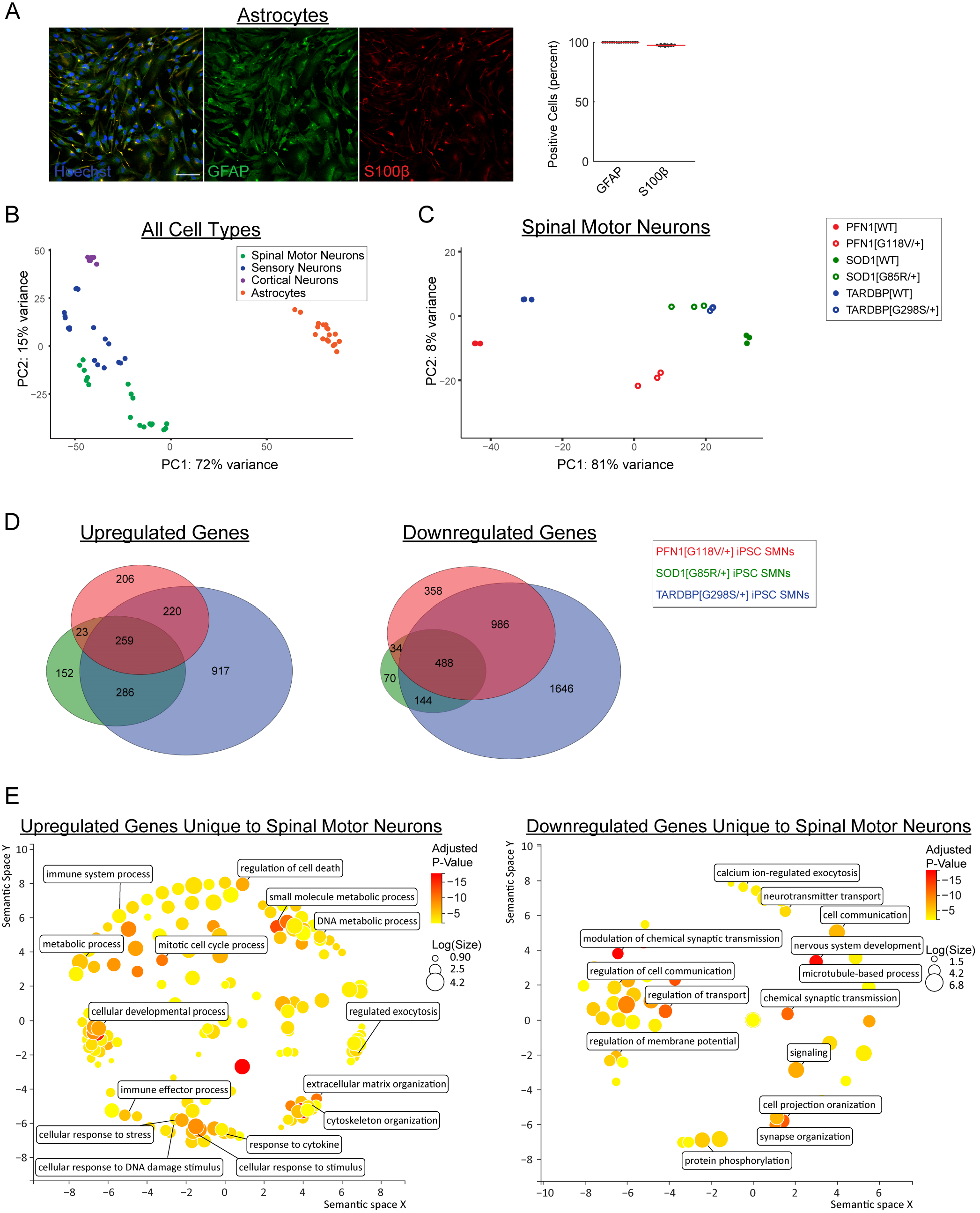
Additional RNA-seq information. (A) Differentiation of astrocytes and staining for astrocyte markers in the parental line used for iPSC editing (scale bar = 100um; n=16 wells per group). (B) Principal component analysis of all RNA-seq libraries. (C) Principal component analysis of SMN RNA-seq libraries. (D) Venn diagram comparing differentially expressed genes in fALS iPSC SMNs vs all controls. (E) Gene ontology analysis of upregulated and downregulated genes specific to SMNs.

**Figure S3.**
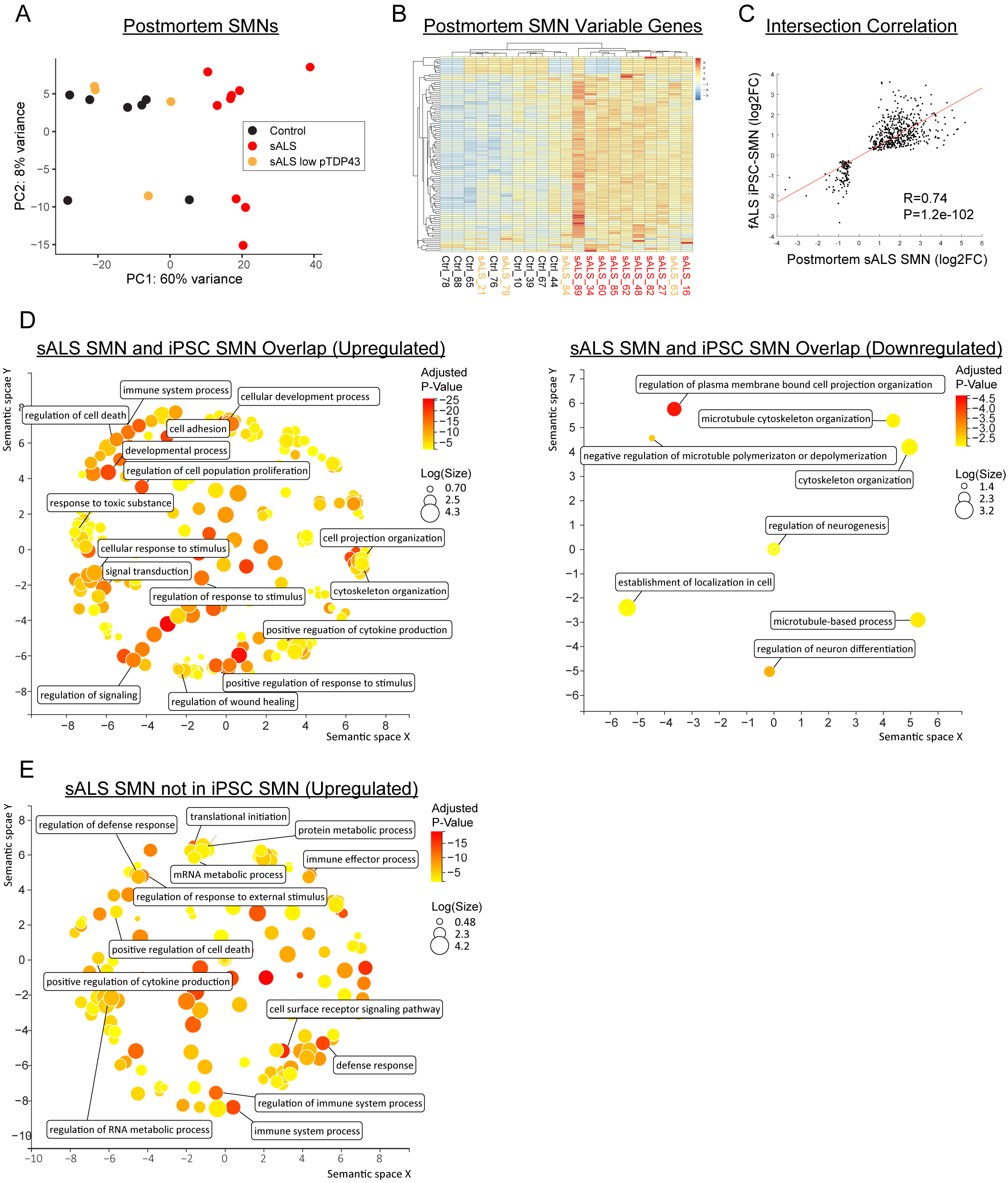
Comparison of differential gene expression in fALS iPSC SMNs (this study) to postmortem sALS SMNs (Krach et al. 2018). (A) Principal component analysis of laser captured postmortem sALS SMNs. (B) Heatmap clustering of genes with the highest variance. (C) Correlation of genes in the intersections of Figure 2F with postmortem sALS SMNs. (D) GO terms for postmortem sALS SMN gene expression changes that are also differentially regulated in iPSC SMNs. (E) GO terms for gene expression changes in postmortem sALS SMNs that are not differentially regulated in iPSC SMNs.

**Figure S4.**
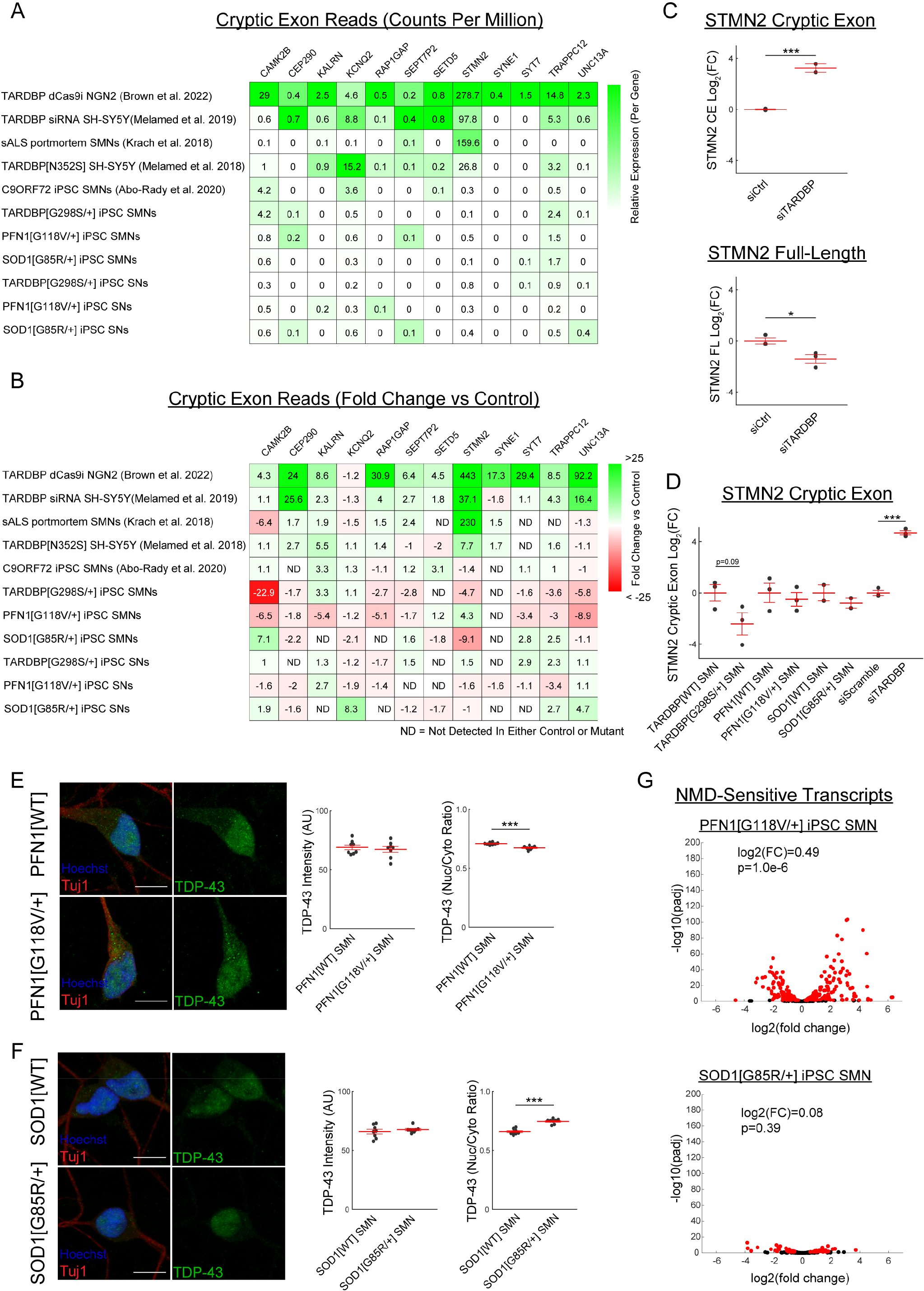
Cryptic exons have very low abundance in iPSC SMNs. (A) RNA-seq counts aligning to cryptic exons in genes with previously reported cryptic splicing. Previously published TDP-43 knockdown experiments have reads aligning to cryptic exons (Brown et al. 2022 and Melamed et al 2019), but neither our iPSC-derived neurons nor another study of *C9ORF72* iPSC SMNs (Abo-Rady et al. 2020) have comparable numbers of reads aligning to cryptic exons despite similar sequencing depths. (B) The same analysis as in (A) but comparing fold change vs control. A pseudocount of 1 was used if reads were detected in only one group. (C) RT-qPCR of *STMN2* cryptic exon and full length when *TARDBP* is knocked down in SH-SY5Y cells, demonstrating fidelity of the primers; n=3 wells for all groups except the *siTARDBP* cryptic exon group where n=2. (D) RT-qPCR using hydrolysis probes for the *STMN2* cryptic exon in iPSC SMN samples and in SH-SY5Y cells with *TARDBP* knockdown; n=3 for all groups except the *SOD1* isogenic pair where n=2. (E) TDP-43 intensity and localization for the *PFN1* isogenic pair and *SOD1* isogenic pair (F); n=8 wells for all groups. (G) Volcano plots of NMD-sensitive transcripts in SMNs with *PFN1^G118V/+^* and *SOD1^G85R/+^* mutations compared to isogenic controls.

